# A ticket to ride - Allele delivery by rail in secondary ruderal colonization by *Arabidopsis arenosa*

**DOI:** 10.1101/171124

**Authors:** Pierre Baduel, Ben Hunter, Sarang Yeola, Kirsten Bomblies

## Abstract

Human-generated ruderal habitats are abundant, but challenging for plants. Some ruderal habitats, however, provide networked corridors (e.g. roadsides and railways) that can facilitate rapid long-distance spread of successfully adapted variants. Here we use transcriptomic and genomic analyses, coupled with genetic mapping and transgenics to understand adaptation to railways in *Arabidopsis arenosa*. We show normally perennial *A*. *arenosa* switched to rapid cycling, a common adaptation for ruderal plants, at least twice upon railway colonization. We further show substantial gene flow from a widely distributed railway colonist likely contributed to secondary colonization by a non-ruderal type, highlighting how connectivity can affect adaptability. We find loss of expression of the reproductive repressor *FLOWERING LOCUS C* (*FLC*) is likely primarily responsible for rapid cycling in the widely distributed railway variant. However, a second railway colonist in the Alps also cycles rapidly, but retains high *FLC*. Some alleles in this population encode non-functional proteins, suggesting *FLC* has started to decay, but most are functional. Instead, this population likely circumvents FLC via a derived allele of *CONSTANS (CO)*, which shows strong evidence of selection in this population. Importantly, we find this CO allele arrived via gene flow from the widespread ruderal, where it was also previously under selection. This suggests ruderal adaptation may have been progressive, perhaps in both cases, with FLC-circumvention arising first, and FLC loss arising later but ultimately obscuring its earlier circumvention. These snapshots of railway adaptation highlight that gene flow from widespread ruderals can provide opportunities for subsequent adaptation by local genotypes.

## Introduction

Human-associated ruderal sites, such as railways, roadsides and field margins are challenging habitats for most plants and thus serve as model systems for adaptation. Colonists of such sites must be able to withstand or evade a variety of stresses including high light, temperature fluctuations, late summer droughts, or even human-mediated interventions such as herbicide applications. Rapid cycling, often coupled with loss of perenniality is a common mechanism by which plants can escape drought and other seasonal stresses that are commonly encountered on ruderal sites and has evolved repeatedly (e.g.^1–7^). An important additional factor on railways may also be that rail beds are cleared of plant life in summers, often annually. For example, since about 1920 German railway ballast has been regularly subjected to thermal treatments or herbicide applications at a rate about six times that used in agricultural settings^8^. Such lethal factors provide truncation selection, which can drive rapid trait evolution^9^ and has been suggested as a driver of repeated evolution of rapid cycling in plants inhabiting ruderal and other marginal habitats (e.g.^1,10^). But ruderal adaptation, once attained, can also provide new opportunities. For example, human-generated “corridor” habitats like railways and roads can facilitate rapid long-distance spread of adapted genotypes (e.g. ^11–14^), which can also provide an opportunity for colonists to come into contact with and perhaps hybridize with related populations they would otherwise have been isolated from. Because of the challenges they face, and the potential they have for rapid long distance dispersal, ruderal plants provide models both for adaptation, and the potential role gene flow may play in adaptation.

Here we study the genetic basis of the acquisition of rapid cycling in a widespread railway colonist of *Arabidopsis arenosa*, a close relative of *A*. *thaliana*^15,16^ that exists in both diploid and autotetraploid forms^17^. Most diploid and autotetraploid populations of *A*. *arenosa* are perennial and found on sheltered rock outcrops or slopes usually in forests or on mountains, but within the autotetraploids, one genetic lineage colonized lowland ruderal sites and is now widely distributed across the railways of central and northern Europe^18^. All railway plants tested to date are rapid cycling and lack a vernalization response (need for winter cold exposure), in contrast to their relatives in mountain sites, which are all perennial and late flowering in the lab^19^, albeit to varying extents.

We showed previously that a railway population that is early flowering in the lab has lost expression of a core floral repressor called *FLOWERING LOCUS C* (*FLC*)^19^. In *A*. *thaliana FLC* expression directly delays flowering^20^, but becomes epigenetically silenced by prolonged exposure to cold (vernalization). Once *FLC* is silenced (or lost by mutation), the repression of flowering promoting genes such as *FLOWERING LOCUS T* (*FT*) and *SUPPRESSOR OF CONSTANS 1* (*SOC1*) is alleviated, allowing them to promote the transition to reproductive development^21^. Our prior observation that an early-flowering railway population lost *FLC* activity suggests parallel evolution between *A*. *arenosa* and related species, as *FLC* has been lost repeatedly to cause early flowering in *A*. *thaliana* accessions^22–29^, and is also associated with a switch to rapid cycling and loss of perenniality in *Arabis alpina*^30,31^. These first observations led us to hypothesize that rapid cycling arose once in *A*. *arenosa* via loss of *FLC*, and that the railway colonists, armed with this adaptation, then spread across the European rail network after it became widely connected in the late 1800’s. In this study, we show a more complex picture was underlying the colonization process.

Here we study the cause underlying early flowering in a widespread railway lineage, as well as an apparently independent railway colonist from the railway at Berchtesgaden in the Bavarian Alps. Plants from this latter population are early and perpetually flowering like other railway populations, but are genetically more similar to plants sampled from mountain populations in the Alps, suggesting an independent colonization of railways from a mountain background and perhaps independent acquisition of early flowering^18^. We show that this secondary colonization was accompanied by gene flow from the more widespread flatland railway populations and that some introgressed loci came under selection in the mountain railway. We use transcriptome analyses, genomic scans for selection, transgenic tests, and genetic mapping to explore the molecular basis of variation in the flowering response in *A*. *arenosa*, and to ask whether introgression from the widespread railway lineage contributed to local flowering time adaptation in a secondary mountain colonist. Our work thus not only provides insight into the mechanisms of ruderal adaptation and loss of perenniality, but also underscores the important role that widely connected ruderals can play, via gene flow, in secondary colonization by additional genotypes.

## RESULTS

### BGS, a transcriptomic outlier among railway populations of *A*. *arenosa*

To study the mechanism(s) of ruderal adaptation in *A*. *arenosa*, we first we sought to identify genes whose expression is correlated with flowering time. To do this, we grew plants from seeds collected from three exposed railway sites (TBG, STE, and BGS) and four sheltered hill / mountain populations (SWA, HO, KA, CA2; Fig 1A; Table S1). In laboratory conditions, railway plants were almost all early flowering and not vernalization (prolonged cold treatment) responsive, while mountain plants show wider variation, but are consistently later flowering and respond strongly to vernalization (Baduel et al.^19^; Fig. 1B). We quantified gene expression in leaves of three 3-week-old unvernalized individuals from each population using read counts from whole-transcriptome sequencing (RNA-seq) aligned to the closely related *A*. *lyrata* reference genome^32^. Principal Component Analysis (PCA) of the genome-wide transcriptional profiles groups early flowering plants from geographically distant railway populations TBG (SW Germany) and STE (Central Poland) very closely. However, the equally early BGS railway population from the Alps (SE Germany) grouped more closely with late flowering mountain populations, albeit in an intermediate position (Fig 1C), suggesting it might be an independent colonist.

**Figure 1.**
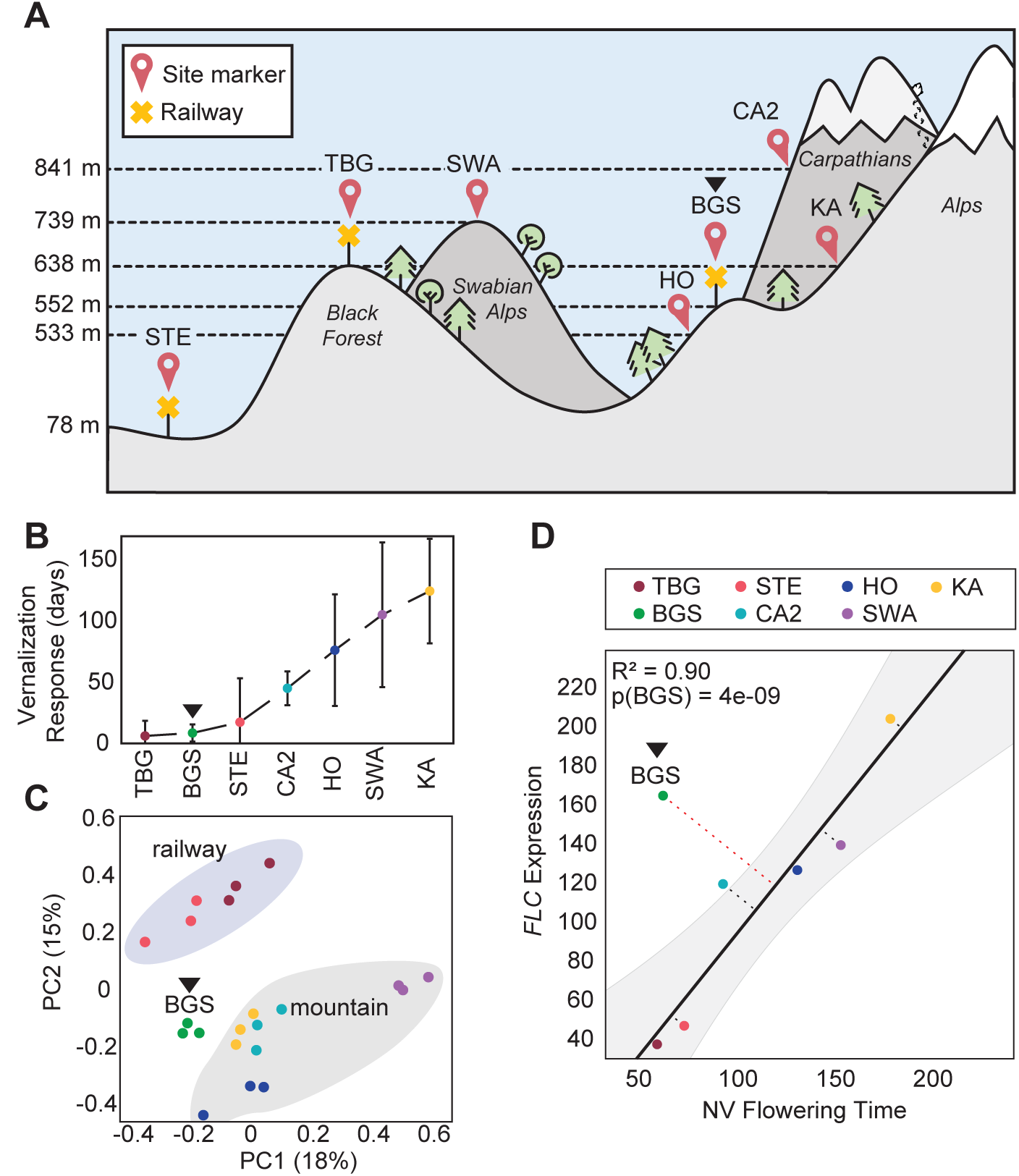
BGS, a transcriptomic outlier among railway populations. (A) Schematic representation of the habitats and altitudes where the railway populations (yellow cross markers): TBG, STE, and BGS (black triangle across all subfigures) and mountain populations: SWA, KA, HO, CA2 were sampled (GPS coordinates in Table S1). (B) Vernalization response of populations reproduced from Baduel et al. as the difference between non-vernalized and vernalized flowering time. All railway populations present almost identical null vernalization responses. (C) First two principal components (PC1 and 2 with percentage of variance explained) of Principal Component Analysis (PCA) of the expression profiles of the 500-most variable genes. Railway and mountain populations group closely together by site-type. (D) Correlation analysis between *FLC* average expression and average non-vernalized (NV) flowering time. The linear regression model after exclusion of BGS is plotted in solid black. Grey area represents the 95% predicted confidence intervals around regression line. Dotted lines are the residual (orthogonal distance) for each data point from the regression line. The p-value for the likelihood to obtain a residual as observed with BGS from residual distribution is indicated as p(BGS).

We next asked which genes are correlated with flowering time variation across our *A*. *arenosa* samples. We identified 76 “flowering-correlated” genes (Table S2, see Methods). These genes are functionally diverse and include only one known flowering time gene^33^, the floral repressor *FLC*. Expression of *FLC* is strongly correlated with flowering time in wild accessions of *A*. *thaliana*^22^–^29^ and here we found a general trend of low *FLC* expression in early-flowering and high *FLC* in late-flowering accessions. The latest flowering plants (from the mountain site, KA) have the highest *FLC* expression while two early flowering railway populations (TBG and STE) have virtually undetectable *FLC* (Fig. 1D), thus putting *FLC* among the 5% most strongly differentially expressed genes between railway and mountain accessions (Fig. S1). The mountain railway population BGS, however, was a striking outlier: When excluding BGS, *FLC* expression levels were very strongly correlated with flowering time (R^2^ = 0.90; Fig. 1D), but though they flower as early as plants from STE and TBG (Fig. 1B), BGS plants have high expression levels of *FLC* comparable to the latest flowering mountain populations (Fig. 1D). Other genes among the flowering-correlated genes show a similar trend: BGS shows expression levels characteristic of early flowering plants for only 18 of the 76 flowering associated genes (Table S2).

The findings for flowering-correlated genes prompted us to ask how much other genes in the genome reflect “mountain-like” expression in the ruderal BGS plants. To more quantitatively assign genes genome-wide as having railway-like or mountain-like expression in BGS, we built a simple metric we called RW/MT_*stat*_ (see Methods). RW/MT_*stat*_ is positive when expression of a gene in BGS is closer to that seen in railway populations and negative when BGS levels are more similar to mountains. The distribution of RW/MT_*stat*_ among differentially expressed (DE) genes was heavily shifted toward negative values, confirming that the gene expression profile of BGS is overall closer to mountain than railway (Fig. 2A). We then selected genes where the RW/MT_*stat*_ values were more extreme than the two-tailed genome-wide 5% thresholds and obtained 872 genes differentially expressed between mountain and railway populations with either railway-like (239 genes) or mountain-like (633 genes) expression in BGS (Fig. 2A). Among these were four flowering-time genes (based on flowering-time list^33^): *FLC*, *VIP5*, and *SSR1* had mountain-like expression in BGS, while *SUPPRESSOR OF OVEREXPRESSION OF CONSTANS1 (SOC1)* had railway-like high expression (Fig. 2A & 2C). *SOC1* promotes flowering, and is directly repressed by a complex of FLC and another protein, SVP in *A*. *thaliana^34,35^* (the gene encoding SVP is also highly expressed in BGS; Fig. 2B). *SPL4*, a direct target of *SOC1*^36^,is also expressed higher in BGS than in late flowering mountain plants (Fig. 2E).

**Figure 2.**
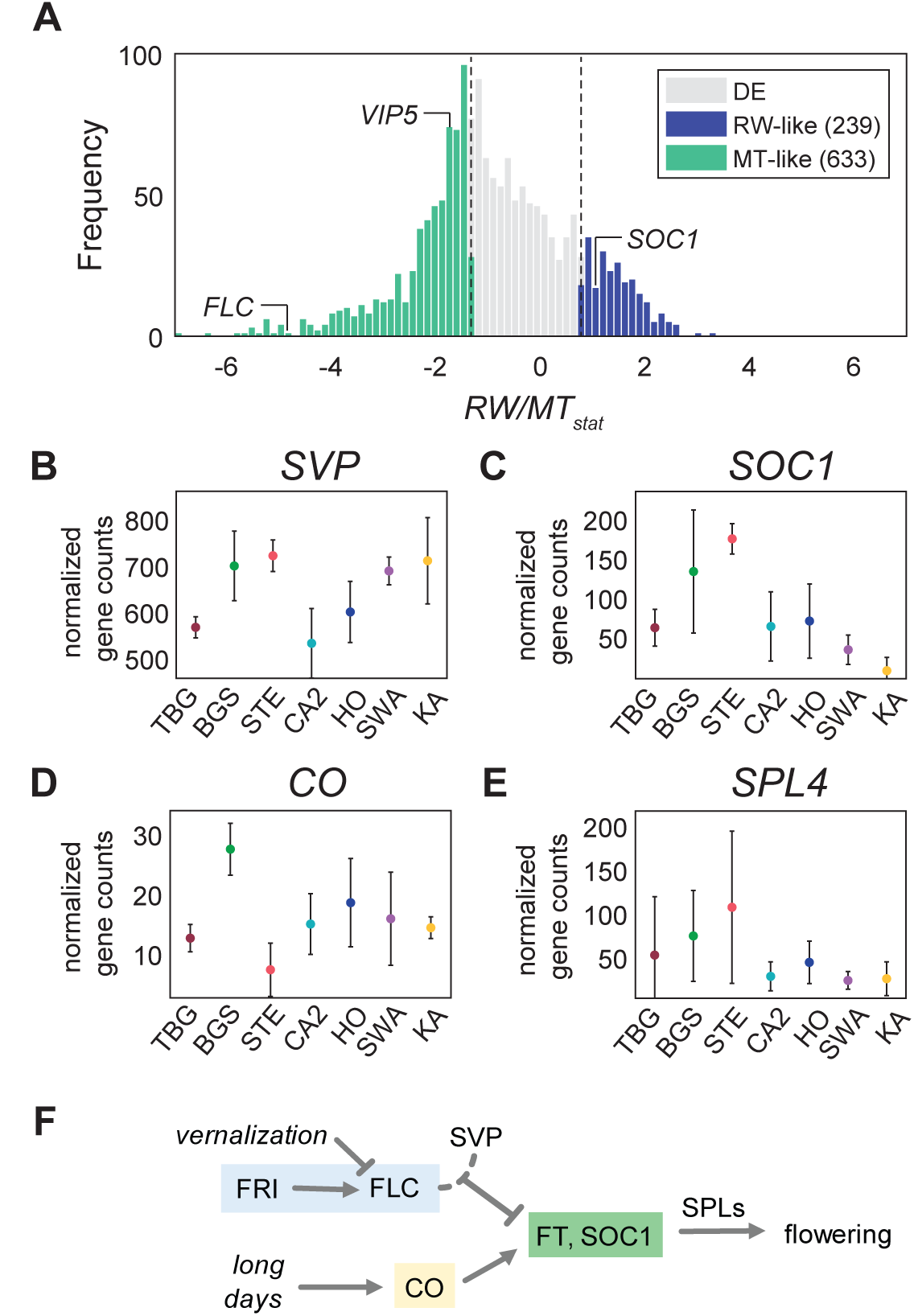
Gene expression patterns in BGS. (A) Railway (RW) and mountain (MT) like expression patterns in BGS measured by RW/MT_*stat*_ among genes differentially expressed (DE) between mountain and railway populations. (B, C, D, E) Normalized expression levels of *SVP* (B), *SOC1* (C), *CO* (D), and *SPL4* (E) across populations (error bars: SD). (F) Schematic representation of the interaction between vernalization (blue) and photoperiod (yellow) pathways. On one side the vernalization pathway represses the expression of flowering activators *FT* and *SOC1* through the FLC-SVP complex, while on the other the photoperiod pathway integrator *CO* activates them. Among the cascade of downstream targets of *SOC1* and *CO* are *SPL* factors including, *SPL4*^21^.

The railway-like elevated *SOC1* and *SPL4* expression in BGS (Fig. 2C) is consistent with its early flowering, but its high expression of *FLC* is not. This suggests that in these plants either *FLC* is not effective in repressing *SOC1*, or *SOC1* activation occurs despite high *FLC* (i.e. that *FLC* is active, but circumvented). A plausible candidate for such circumvention is *CONSTANS* (CO), which is a direct activator of *SOC1* and can activate it even in the presence of high *FLC* levels^37,38^ (Fig. 2F). We observed variation in *CO* expression levels, with BGS having the highest levels, but overall normalized gene counts remained low (below 30) for all populations including BGS (Fig. 2D).

### Indications of *FLC* decay in BGS

Since high *FLC* expression is generally associated with late flowering in both *A*. *thaliana*^22^, and *A*. *arenosa* (this study), we hypothesized that the allele expressed in BGS might be non-functional. In *A*. *arenosa* the *FLC* locus contains two full-length (*AaFLC1*, *AaFLC2*) and one truncated (*AaFLC3*) copy^39^, so we first established which copies are expressed using an approach we used previously (Baduel et al.^19^). We found that 90% of *FLC* expression was contributed by *AaFLC1* in all mountain populations, as well as BGS, while *AaFLC2* contributed the remaining 10% (Fig. S2); *AaFLC3* expression was undetectable in any population, and thus we did not study it further. From our genomic sequence data, we found no changes in *AaFLC2* in BGS relative to late flowering mountain populations, but one allele of *AaFLC1* in BGS has two non-synonymous derived polymorphisms in exons 3 and 4 that are unique to BGS, where they are found at frequencies of 0.22 and 0.19. These two polymorphisms were usually within the same haplotype (in 6 out of 7 PCR clones). The more frequent allele is identical to that found in late-flowering KA, and thus the encoded protein is likely functionally identical to that in KA and we did not study it separately.

To test whether the rarer *FLC* variant in BGS is functional, we isolated a cDNA of the BGS *AaFLC1* allele with the two amino acid changes, *AaFLC2* from BGS, and *AaFLC1* and *AaFLC2* from late-flowering KA. We expressed these under a constitutive promoter (35S) in the early flowering *A*. *thaliana flc*-*3* mutant (Fig. 3). We phenotyped over 30 independent transgenic lines for each of the four constructs for their leaf number at bolting (LNB), a commonly used measure of flowering time in *A*. *thaliana*. LNB was higher than the maximum observed in the Col-0 *flc*-*3* line (LNB = 13) for 23% of lines expressing KA *AaFLC1* and *AaFLC2* (Fig. 3; 14 out of 62 and 24 out of 105 respectively). 16% of transgenic lines carrying *AaFLC2* from BGS were also had higher LNB than *flc*-*3* alone (9 out of 57), but only 5.7% of lines with BGS *AaFLC1* did (2 out of 35) (Fig. 3). To ask whether this variation was likely due to FLC, we measured *FLC* expression by qRT-PCR in each of the transgenic lines. For *AaFLC1* and *AaFLC2* from KA and *AaFLC2* from BGS, there was a good correlation between *AaFLC* expression level and flowering time, suggesting these genes encode active versions of *FLC* (R^2^ > 0.4; Fig. S3). For BGS *AaFLC1*, however, there was no correlation between transgene expression and flowering time (R^2^ = 0.02). The two late flowering lines had low or null *AaFLC1* expression while lines with high *AaFLC1* expression were not late-flowering. This lack of correlation suggests that the variant BGS *AaFLC1* allele is not functional in *A*. *thaliana*, at least in terms of floral repression.

**Figure 3.**
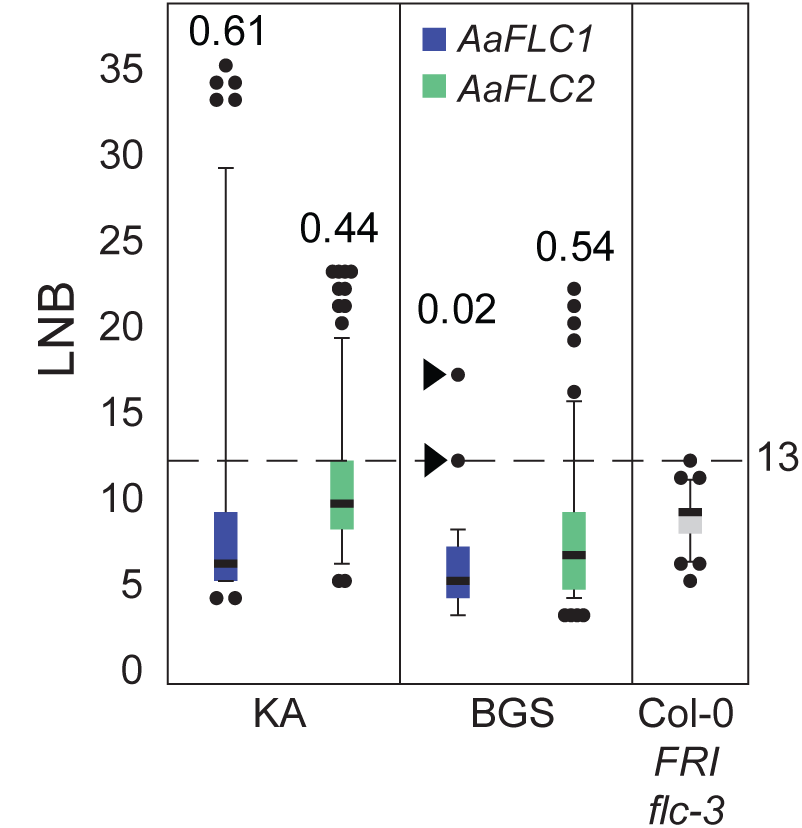
Non-functional *AaFLC1* BGS allele in transgenic *A*. *thaliana*. Boxplot of flowering time measured by leaf number at bolting (LNB) of Kanamycin-resistant T1s with 35S-driven cDNAs of both *AaFLC1* (blue) and *AaFLC2* (green) from KA (left panel) and BGS (middle panel). For comparison the flowering time of the FRI flc-3 Col-0 background line is plotted in the third panel in grey. The latest flowering time observed in the background line (LNB=13) is represented by a dotted line. The numbers above each box indicate the correlations between LNB and *FLC* expression (Fig. S3). Two late-flowering transgenic individuals obtained with BGS *AaFLC1* that nevertheless have low FLC expression are indicated with black triangles.

### A mountain genotype with introgression of railway alleles

Our PCA results (Fig. 1C) and our previous biogeography work^18^, suggest that BGS is primarily a mountain genotype, but has extensive shared polymorphism with the widespread flatland railway type (which is not true of other mountain populations^18^). To analyze the extent and patterns of this shared diversity we complemented previously-generated whole-genome sequences^19,40,41^ with additional sequencing to reach a total of 47 individuals from 2 railway (TBG and STE) and 3 mountain populations (HO, GU, and KA) and BGS. PCA of the genome sequence data recapitulated the pattern observed with the transcriptome: despite their geographic separation, flatland railway populations TBG and STE grouped tightly together and were clearly separable from mountain populations, while BGS was again intermediate but closer to the mountains (Fig. 4A). To further analyze population structure of our samples, we used STRUCTURE^42^ on 627,016 SNPs. The ΔK ad-hoc statistics^43^ support that our populations form 2 major clades comprising a railway clade including TBG and STE, and a mountain clade of HO, GU, and KA (Fig. 4B). The 8 BGS individuals clearly showed a hybrid genomic constitution between the two groups.

**Figure 4.**
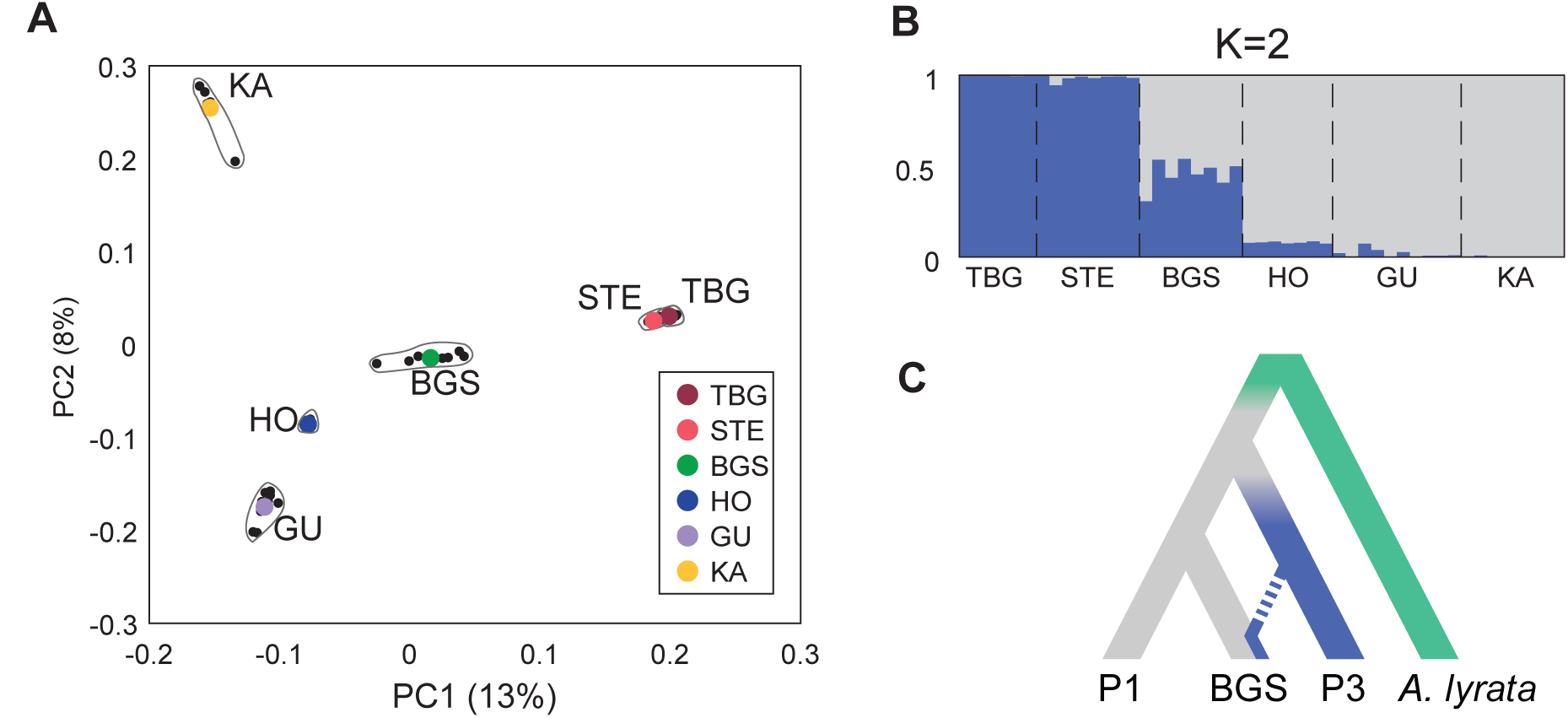
BGS shares genomic variation with railway and mountain backgrounds. (A) First two principal components (PC1 and 2 with percentage of variance explained) of Principal Component Analysis (PCA) of the genomes from 47 re-sequenced individuals of three railway (TBG, STE, BGS) and three mountain (HO, GU, KA) populations. (B) Genomic clustering of individuals using STRUCTURE with K=2. Each individual is represented by a single vertical line broken into K=2 segment with length of each colored bar proportional to the posterior probability of belonging to each cluster. (C) Population history model used for ABBA-BABA evaluation of introgression fraction within BGS, with background and donor populations P1 and P3 and *A*. *lyrata* used as outgroup.

We quantified the genome-wide fraction of introgression with the modified *f*-statistic *f̂_hom_* described by Martin et al.^44^. Using *A*. *lyrata* as a reference, we used either mountain or railway populations as the donor population (P3) or the background population (P1) in the ABBA-BABA configuration (respectively columns and rows of Table 1, represented schematically in Fig. 4C). We excluded GU because it has experienced gene flow from *A*. *lyrata*^41^, which would bias estimates of introgression. When we used the mountain populations HO or KA as donors (P3), the estimates of introgression into BGS were ~20% higher than when we used these as background (P1), consistent with BGS being overall more mountain-like. If we thus assume BGS has a mountain origin, the fraction of introgression from the flatland railways was estimated between 8.7% and 11.9% (Table 1).

**Table 1.** Fraction of introgression *f̂_hom_* with +/- giving jackknife standard deviation

|  |  | donor (P3) |  |  |  |
| --- | --- | --- | --- | --- | --- |
| | $\hat{f}_{hom}$ | TBG | STE | HO | KA |
| background (P1) | TBG | | | 30.40%<br>(± 0.5%) | 26.26%<br>(± 0.41%) |
| | STE | | | 33.03%<br>(± 0.48%) | 28.12%<br>(± 0.37%) |
| | HO | 9.75%<br>(± 0.4%) | 11.91%<br>(± 0.63%) | | |
| | KA | 8.67%<br>(± 0.33%) | 10.53%<br>(± 0.51%) | | |

We then used another metric for the fraction of introgression (*f̂_d_*) to identify introgressed loci on a finer scale. *f̂_d_* performs better than Patterson’s D and *f̂_hom_* for identifying introgression on a small window basis^44^. For each mountain-railway (P1-P3) couple we calculated per gene estimates of *f̂_d_* for any genes with more than 25 informative SNPs. We then scanned for genes with a high fraction of introgression (*f̂_d_* > 3 ∗ *f̂_hom_*) and kept only loci presenting high *f̂_d_* in all 4 comparisons. This way we identified 1180 candidate introgression loci from flatland railways into BGS (Table S3). We next asked if genomic regions introgressed from railways into BGS had more railway-like expression profiles. Among the 1180 genes putatively introgressed from other railway populations into BGS, only 53 (5%) were differentially expressed between railway and mountain populations. Of these, 11 (21%) have railway-like expression in BGS and 14 (26%) had mountain-like expression in BGS (Table S4), showing that both cis and trans effects occur, and that overall there is no clear trend whether introgressed loci reflect the expression levels characteristic of the donor or the recipient.

### Genetic mapping of Flowering Time

To assess whether the introgressed genes or DE genes we identified might be responsible for the early flowering phenotype of the mountain and flatland railway populations, we took a genetic mapping approach. We grew 795 and 845 F_2_ plants derived from TBG × SWA and BGS × KA crosses respectively (both SWA and KA are late flowering mountain types) and quantified flowering by time to bolting (initiation of inflorescence outgrowth). The phenotype distribution of TBG × SWA F_2_ plants showed three modes, around 42, 55, and 68 days to bolting with a tail of plants that did not bolt before the end of the experiment at 80 days (approximately 1/8 of plants; Fig. S4). In contrast, the distribution of BGS × KA F_2_ plants was bi-modal with a single early-flowering mode around 50 days and a group of very late flowering individuals (~1/4 of plants) that did not bolt before the end of the experiment, despite extension to 140 days. The difference between the two phenotype distributions suggests a distinct genetic basis for early flowering in TBG × SWA vs BGS × KA. As these *A*. *arenosa* populations are autotetraploids with tetrasomic inheritance^40^, a ¼ proportion of late plants in BGS × KA could be explained by segregation of a dominant locus that was present in a single copy in each of the two F_1_ parents (i.e. both had genotypes Aaaa, where the A allele confers early flowering).

We used a restriction-associated reduced representation sequencing (RADseq) approach to genotype 452 and 284 individuals selected from each mode in the two populations. We obtained 7907 and 8666 informative single nucleotide polymorphisms (SNPs) for the two populations after filtering for a minimum coverage of 40% of individuals per SNP, and tested for correlations of markers in each group with flowering time. Surprisingly, in both populations there was a very strong association with days to bolting in overlapping regions on the upper arm of scaffold 6 and nowhere else in the genome (Fig. 5). Regions of high LOD scores (>30) in TBG × SWA and BGS × KA spanned 6.5 Mb and 6.8 Mb respectively, with a shift down the chromosome in BGS × KA (Fig. 5). The peaks are strongly significant in both F_2_ populations (p-values of marker with highest LOD: 2e-18 in TBG × SWA and 4e-13 in BGS × KA). Together the two intervals of high LOD in the two populations define a 7.8 Mb region hereafter referred to as FT-peak, which spans 1877 genes. In both TBG × SWA and BGS × KA the SNP of maximum LOD score falls close to *FLC* (within 135kb and 75kb respectively; Fig. 5), though other genes known to be associated with flowering time in *A*. *thaliana* are also found within the FT-peak region, including *MYB33* and *CO*.

**Figure 5.**
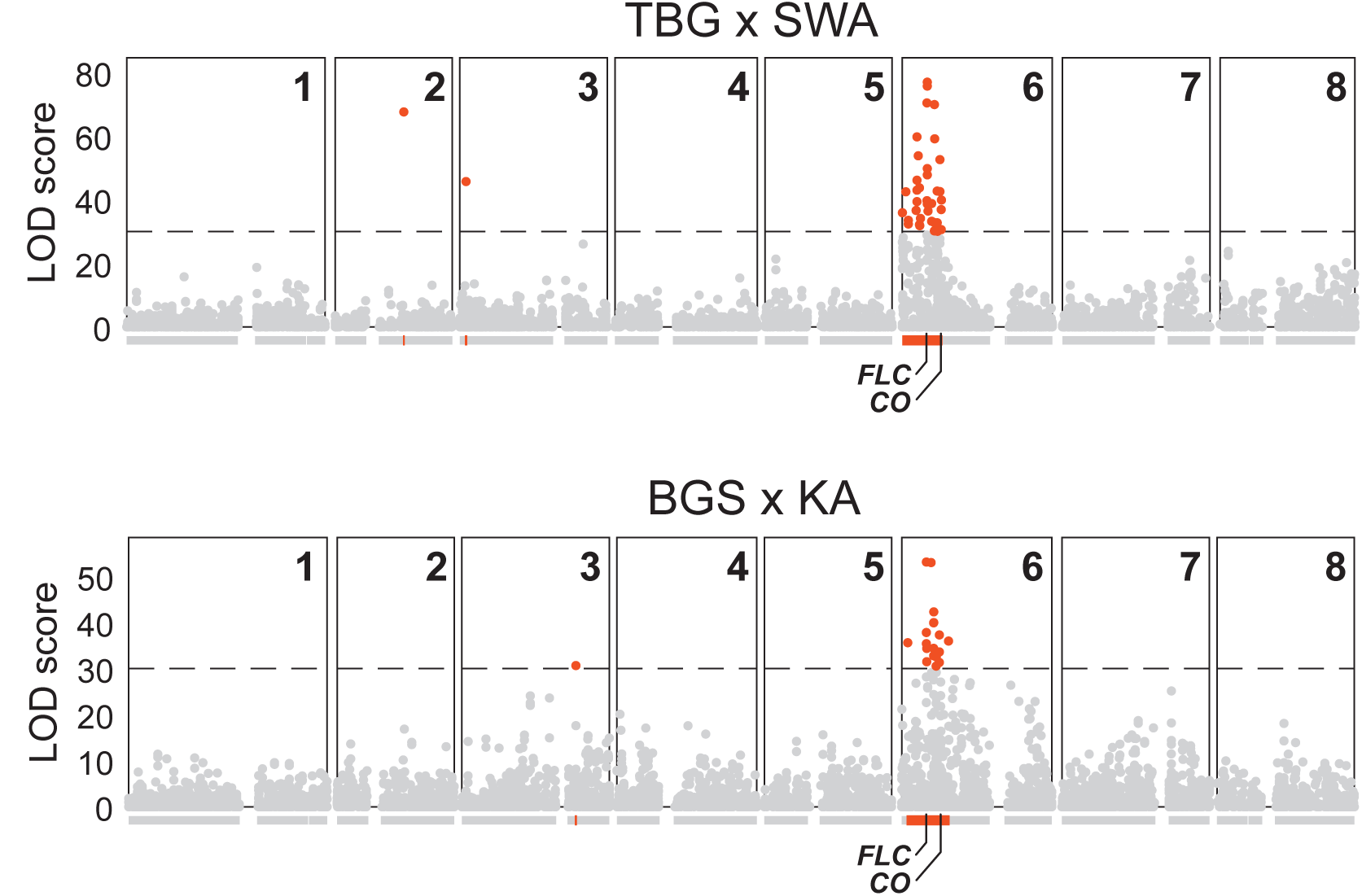
BSA mapping of flowering time in TBG × SWA and BGS × KA F_2_s. Distribution of single-marker LOD scores for time to bolting across the 8 scaffolds in TBG × SWA and BGS × KA F_2_s respectively above the gene models for each scaffold. High LOD (LOD>30) markers and genes within high-LOD regions are highlighted in red.

Among the 5% most strongly differentially expressed genes between early and late flowering plants within the FT-peak region, *FLC* was the only one among 174 *A*. *thaliana* genes previously associated with flowering time regulation^33^ (Fig. S5B). Though not significantly differentially expressed, *CO* is an intriguing candidate for the BGS × KA cross, because in *A*. *thaliana* high CO activity can bypass the repression of reproduction caused by high *FLC* expression in a dominant fashion^38^. In TBG × SWA, *CO* is located at the very edge of the interval within 78kb of the most downstream SNP of LOD score above 30, but in BGS × KA *CO* is located over 1.3 Mb from the end of the high-LOD region (Fig. 5). We also estimated individual effects of high LOD markers across the FT-peak region (see Supplemental Text 1) and this supported the potential involvement of both *FLC* and *CO* in early flowering in the BGS × KA cross.

### A derived railway-specific *CO* haplotype under positive selection

We next investigated whether any of the regions introgressed into BGS from other railway populations subsequently came under selection in BGS using Fay & Wu’s H, a statistic sensitive to excess of high-frequency variants compared to neutral expectations^45^. We found 128 genes among the 1180 candidate introgressed regions where windows with Fay & Wu’s H values are within the most extreme 5% negative outliers genome-wide. Among these 128 genes is *CO*, which is the only gene on this list with a known role in regulating flowering time (based on list by Fornara et al.^33^). We then asked if any of these genes might have already been under selection in the source railway populations by scanning for windows with high genetic differentiation between railway and mountain groups (top 5% windows for **G**_ST_, which is F**ST** generalized to multi-allelic sites^46^; Fig. 6A, Fig. S6A). We considered only windows that were also outliers for Fay & Wu’s H in both TBG and STE, but not in the mountain populations. By these criteria, 24 genes had marks suggesting railway-specific selection (Table S5). Three of these genes were also outliers for Tajima’s D, which is sensitive to scarcity of low-frequency variants, a complementary mark of positive selection^47^. *CO* fulfilled all of these criteria (Fig. 6B, Fig. S6B).

**Figure 6.**
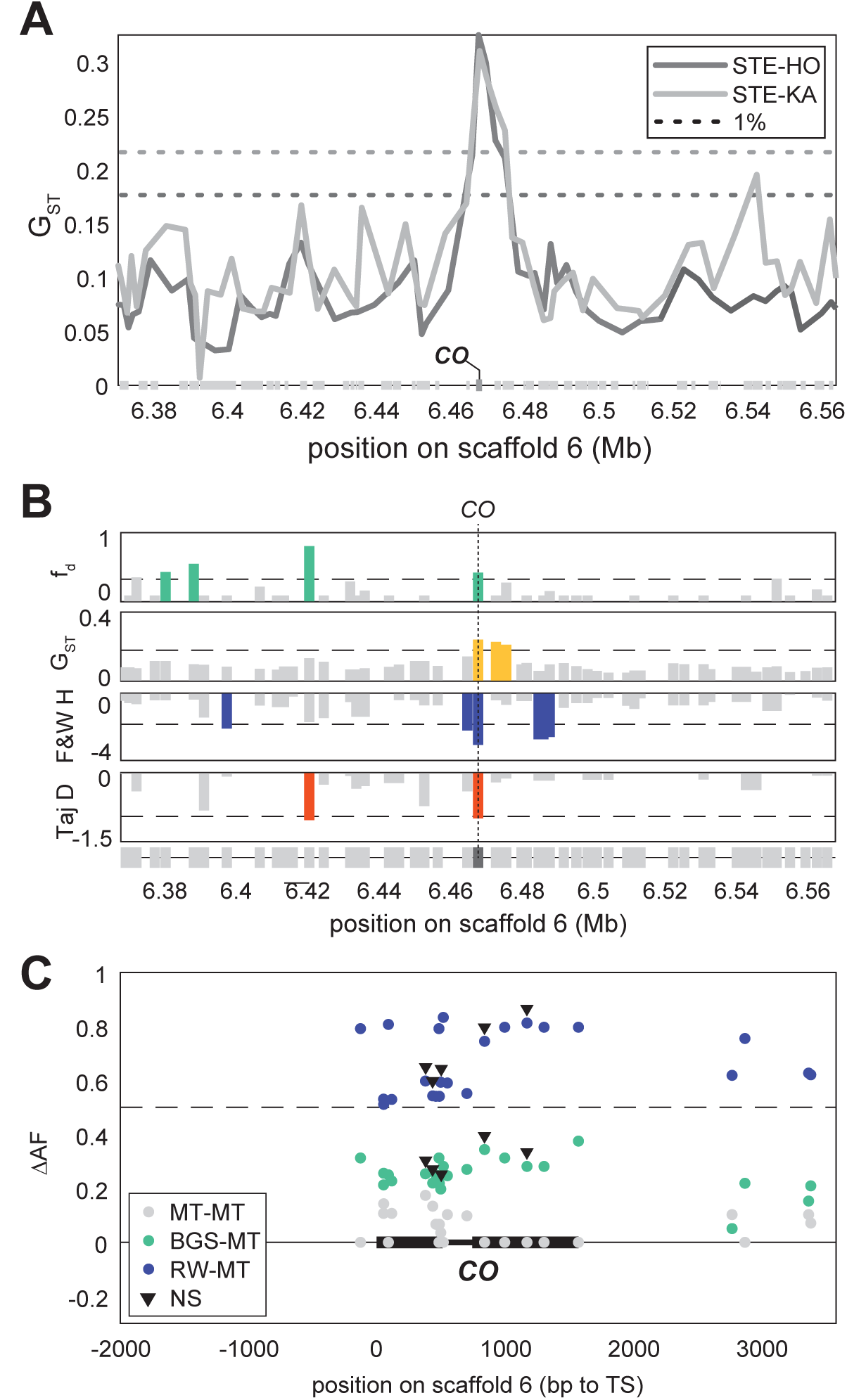
Railway-specific selection on a highly-differentiated railway haplotype of *CONSTANS*. (A) Marks of differentiation between one railway population (STE) and two mountain populations (HO and KA) evaluated with G_ST_ across *CO* region. Dotted lines are respective genome-wide 1% threshold levels. (B) Gene-wise marks of introgression (*f̂_d_*), railway-mountain differentiation (G_ST_), and railway-specific positive selection (Fay & Wu’s H, and Tajima’s D) across *CO* region. For each gene, only the least extreme values are represented. Dotted lines are 3 ∗ *f̂_d_*(*KA*,*BGS*,*TBG*) (upper panel) or most extreme genome-wide 5% threshold levels. (C) Average allele-frequency differences (ΔAF) of 23 SNPs with ΔAF(RW-MT)>0.5 (dotted line) between railways and mountains (RW-MT in blue), BGS and mountains (BGS-MT in green) and HO and KA (MT-MT in gray) around CO. Non-synonymous (NS) mutations are indicated with black triangles.

*CO* shows a clear pattern of differentiation between mountain and railway populations (Fig. 6A). Within 200bp of the coding sequence and across the first exon, we found 23 derived polymorphisms (relative to the reference *A*. *lyrata* genome^32^) at frequencies more than 0.5 higher in railway populations than in mountains. Ten of these are unique to railway populations. These polymorphisms segregate at frequencies averaging 0.79 (sd = 0.03) in both STE and TBG, and are absent from all three mountain populations sampled. Five of the 23 high-frequency variants in the first exon of *CO* are predicted to cause non-synonymous amino-acid substitutions, including two of the SNPs unique to railways (Fig. 6C). All 10 polymorphisms were also found in BGS at frequencies around 0.30 (sd = 0.04). At these frequencies, if CO has a dominant effect (e.g. by circumventing *FLC* expression^38^), 99.8% of STE plants and 76% of BGS plants would have the relevant early-flowering phenotype (calculated as 1-q^4^ where q=0.7 and represents the frequency of the ancestral allele). This estimate assumes Hardy-Weinberg equilibrium and tetrasomic inheritance, which we have previously shown to be valid for *A*. *arenosa^40^*.

## DISCUSSION

### Gene flow between a widespread ruderal and a new colonist

In contrast to mountain populations, most railway populations within autotetraploid *A*. *arenosa* form a single genetic lineage spread over hundreds of kilometers across central Europe, consistent with the idea that railways and roadsides provide “corridor” habitats that can facilitate rapid dispersal of adapted colonists^11-14^. However, one population we sampled from a mountain railway site in Berchtesgaden, Germany (BGS), appears to be a hybrid primarily carrying mountain genotypes. Our data suggest BGS is likely an independent colonist from the mountains that sustained substantial gene influx from previously existing flatland railway populations. We cannot rule out, however, that it might not have gone the other way – namely that BGS might have been first colonized by a railway type, that then was genetically “swamped” by gene flow from adjacent mountain populations. In either case, however, the genes that have become, or remain, distinctly railway-like in this population seem to be the products of selection acting to favor alleles of flatland railway origin in an otherwise mountain genome. This highlights the potential for introgression of alleles from widespread railway ruderals into neighboring non-ruderal populations.

The rail networks in Germany and Poland became widely connected in the mid to late 1800’s (Fig. S7), but the widespread “railway lineage” seems to have diverged from other *A*. *arenosa* earlier than that^18^, suggesting it inhabited a similar habitat elsewhere (e.g. mountain scree slopes or river cobbles) that allowed it to rapidly colonize railways as the networks were built. Subsequent spread of *A*. *arenosa* along railways thus allowed contact between genotypes that were previously geographically isolated (as most modern mountain *A*. *arenosa* genotypes still are^41,48^) and thus the railway lineage acquired (with inadvertent human assistance) the potential to act as a conduit of gene flow. Colonization of the Berchtesgaden railway was likely much more recent. The railway to Berchtesgaden was built in 1888, and completely rebuilt in 1940 to accommodate sudden heavy traffic to Hitler’s infamous Eagle’s Nest, built above Berchtesgaden in 1937. BGS is primarily mountain-like, but has sustained substantial gene flow from flatland railway plants. Importantly, this includes several potentially adaptive genes that show marks of having been under selection after introgression, including a flowering time gene (*CO*) that is a good candidate for driving early flowering in BGS despite its high expression of *FLC*. This raises the possibility that some adaptive alleles in BGS arrived by gene flow, and is consistent with a growing number of examples of “adaptive introgression”^49–54^ having played a role in local adaptation. Furthermore, these findings highlight that “corridor ruderals” can affect the adaptive process in local populations they come into contact with. On the other hand, this new colonist brings novel alleles from its original mountain home to the railways, and in follow-up work it will be interesting to ask whether adaptive introgression is a two-way street, or whether the benefit of hybridization comes only to the new colonist.

### Genetically distinct early flowering in BGS and other railway populations

Flatland railway *A*. *arenosa* have entirely lost *FLC* expression (this study and Baduel et al.^19^), which likely explains their early flowering. This connection is supported by our mapping data, which showed that markers close to *FLC* were strongly associated with additive effects on flowering time in F_2_ populations. This parallels findings in *A*. *thaliana*, where multiple independent losses of *FLC* are seen in early flowering accessions^22–29^, as well as in *A*. *alpina*, where losses of the *FLC* homolog *PEP1* are similarly associated with a switch from episodic flowering and a requirement for vernalization, to rapid cycling and perpetual flowering^30,31^.

Our initial hypothesis was that the mountain railway population, BGS, which is as early flowering as flatland railway populations, shares the same genetic basis for earliness and acquired this through gene flow from other railway populations. However, this seems to be only partly correct. First, we found that BGS is genetically more mountain-like, and shows more mountain-like gene expression genome-wide. Importantly, this was true for 74% of the 76 genes that we found to be strongly correlated with flowering time in all other *A*. *arenosa* populations, including *FLC*. High expression of *FLC* is usually associated with late flowering^22^, as well as low expression of a key reproductive transition promoting gene, *SOC1*^20^. We found, however, that despite its high *FLC* expression, BGS also has high *SOC1* expression, which could explain its early flowering. How could BGS be early flowering and express high levels of *SOC1* while still having high expression of *FLC?* We considered two scenarios: (1) BGS could be expressing a non-functional allele of *FLC*, and (2) BGS could have somehow circumvented *FLC* activity. We found that one expressed *AaFLC1* allele in BGS is inactive, but is found in BGS at only 20% frequency. The more common allele is identical to the active alleles found in other mountain plants. Thus, while independent *FLC* loss may contribute to early flowering in BGS, it does not seem to be the whole story. Our mapping and gene expression data both support the hypothesis that the mechanisms underlying early flowering in BGS and the flatland railway populations are at least somewhat distinct and that *CO* may be contributing to early flowering in BGS, likely by circumventing *FLC*. *CO* is known from *A*. *thaliana* to be a direct regulator of the flowering promoting gene *SOC1*, which it targets antagonistically with *FLC*^55^. Importantly, high CO activity can circumvent high FLC activity to activate *SOC1* and promote flowering^38^. Thus we believe that CO may be an important factor for early flowering in BGS. Like *FLC*, *CO* has also been implicated in a range of species in natural variation for flowering time, including *A*. *thaliana*, *Brassica nigra*, and rice^56–58^ and may have been a target of selection during rice domestication^59^, suggesting it, too, can be an evolutionary hotspot for flowering time.

In BGS, numerous loci show evidence of introgression from flatland railways, but importantly, several seem to have come under selection after arriving in BGS. One of the strongest signatures of post-hybridization selection is in *CO*, which also shows evidence of having been under previous selection in flatland railway populations. The derived railway allele differs from the mountain allele by numerous tightly linked polymorphisms, several of which cause amino acid changes unique to the railways. If dominant, this allele is found at a frequency that 76% of BGS plants should be early flowering. Thus we hypothesize that although *FLC* may play a role, *CO* may be the primary cause of earliness in BGS, while loss of *FLC* is (currently at least) the primary cause of earliness in the widespread railway lineage. From the observation that the derived *CO* allele shows evidence of also having been under selection in the flatland railways previously, we suspect it may have been the original cause of early flowering in these populations as well. Thus we propose the following hypothesis: that initially after ruderal colonization *FLC* expression in the railway populations would have been high, as it remains in BGS, and that this was circumvented by the novel derived dominant allele of *CO*. Later, as *FLC* became ineffective (because of circumvention by CO), loss of function alleles arose that also contribute to earliness, perhaps with even greater phenotypic effect. The *FLC* loss of function alleles may ultimately contribute more strongly to early flowering than CO. This scenario could also help explain the presence in BGS of an apparent loss of function *FLC* allele at low frequency (about 20%). The observation that at least some introgressed loci in BGS may have come under selection, adds to a growing body of evidence that adaptive introgression may be an important factor in rapid adaptation to challenging environments^41,49-51,60^.

### Summary

In summary, we propose that an initial ruderal adaptation that involved a switch to rapid cycling via *CO* and *FLC* allowed a lineage of *A*. *arenosa* to colonize the extensive rail networks of Europe. This corridor habitat facilitated gene flow and allowed a subsequent colonist from the mountains to establish on railways with the help of gene flow that brought in a derived *CO* allele, along with several other potentially adaptive alleles. This highlights how adaptive gene flow from widespread “weedy” colonists can alter the genetic architecture and adaptive potential of plant species and facilitate gene flow among previously isolated populations.

## MATERIALS AND METHODS

### Plant materials and growth conditions

*A*. *arenosa* seeds used in this study were obtained from the same natural populations described and phenotyped for flowering time by Baduel et al (Table S1). All populations are autotetraploid; either they originate from regions where only autotetraploids occur^61^ or they were confirmed using flow cytometry^40^. We grew sibling arrays from seeds of single individuals growing in nature as previously described^40^ in Conviron MTPC-144 chambers with 8 hours dark at 12°C, 4 hours light (Cool-white fluorescent bulbs) at 18°C, 8 hours light at 20°C, 4 hours light at 18°C. For all plants we recorded germination date by root emergence on agar ½ X Murashige-Skoog medium plates. *A*. *thaliana* plants were grown in similar Conviron MTPC-144 chambers using instead 16 hours light (Cool-white fluorescent bulbs) and 8 hours dark at a constant 22°C.

### Genetic mapping

F2s were generated from both TBG × SWA and BGS × KA parental crosses and phenotyped for flowering time using time to bolting (defined as the time when the inflorescence reached 1 cm tall). For plants that had not flowered by experiment end (80 days for TBG × SWA and 140 days for BGS × KA) we assigned cutoff values. We genotyped individuals in order to cover with at least 130 individuals (~15% of F2s) each tail of the distribution of days to bolting, as well as other phenotypic distributions (days to first open flower, IBN) with which we eventually did not have the statistical power to pursue an informative comparison. We prepared sequencing libraries using a modified double-digest RAD-seq protocol as previously described^18^. Libraries were sequenced on an Illumina HiSeq 2000 with 50 bp paired end reads, to 16x coverage.

We filtered out SNPs not present in a minimum of 40% of individuals, LOD scores and p-values for each SNP were obtained from a simple marker linear regression analysis. For each regression we compared additive, recessive, and dominant models and used the model providing the highest LOD. Within the high-LOD markers (LOD>30) of scaffold 6, we then ran a stepwise multiple linear model (MLM) regression to discard markers not significantly improving the sum of squared errors (F-statistics) after completely discarding individuals with missing data (not supported by MLM regression analysis). Out of this MLM we computed the percentage of variance explained (PVE) of each marker conserved as the semi-partial correlation coefficients which represents the loss of correlation when the marker is taken out of the model. As this method does not take into account the correlated effect of all markers and therefore underestimates the effect of each marker, we compared the semi-partial correlation coefficients with the PVE obtained when simple marker model is applied at each marker. This second method on the other hand overestimate the specific effect of each marker as it does not take into account the effect of surrounding markers.

### RNA isolation, sequencing and analysis

We extracted RNA from leaves of four-weeks-old plants with three biological replicates for each of seven populations (TBG, BGS, STE, KA, CA2, HO, SWA) using the RNeasy Plant Mini Kit (Qiagen). We synthesized single strand cDNA from 500ng of total RNA using VN-anchored poly-T(23) primers with MuLV Reverse Transcriptase (Enzymatics) according to the manufacturer’s recommendations. We made RNAseq libraries using the TruSeq RNA Sample Prep Kit v2 (Illimina) and sequenced libraries on an Illumina HiSeq 2000 with 50bp single-end reads. We sequenced between 9.8 and 18.8 million reads (avg 13.6 million). We aligned reads to the *A*. *lyrata* genome^32^ using TopHat2^62^ and re-aligned unmapped reads using Stampy^63^. We acquired read counts for each of the 32,670 genes using HTseq-count^64^ with *A*. *lyrata* gene models^32^. We assessed quality of the count libraries by PCA and Euclidean distance analysis; biological replicates of each population are most similar to each other (Fig 1C). We normalized for sequencing depth using DEseq2 in R^65^ and further analyses were performed in MATLAB (MathWorks). PCA was performed using the 500 genes with the highest variability, as recommended in the DESeq2 package^65^. The *FLC* locus in *A*. *arenosa* has three tandemly duplicated FLC-like genes^39,66^ of which two, *AaFLC1* and *AaFLC2* are clearly homologous to *FLC* from *A*. *thaliana*. RNAseq reads from the *A*. *arenosa FLC* region do not all differentially map to the two *FLC* duplicates of *A*. *lyrata* so we also aligned transcriptome reads to an updated BAC sequence of the *FLC* region^66^ to better discern read counts for each *FLC* as described in Baduel et al^19^.

Correlations between flowering time and gene expression were calculated between the average non-vernalized flowering time reported in Baduel et al. and the average expression of each gene per population after filtering for genes with normalized expression counts above 10 in at least one sample in order to avoid low expression artefacts. We excluded BGS to obtain the overall correlation coefficient for each gene. We obtained a list of 76 “flowering-correlated” genes (Table S2) by retaining the top 1% most strongly correlated after filtering for genes with significantly different expression between mountain and railway plants (at p < 0.05). We then asked whether the BGS datapoint falls outside the 95% confidence interval of the regression line and calculated how likely its position is given the noise in each trend (as we did for *FLC*, Fig. 1D). For each gene we estimated how likely this BGS residual could be obtained from a distribution of residuals modeled as a normal distribution of mean 0 and sigma estimated as the standard-deviations observed with all other populations (two-tailed comparison). Using these criteria BGS was an outlier for 58 of the 76 flowering-correlated genes, including *FLC*, with mountain-like expression levels for 56 of the 58 outliers (Table S2). The 18 strongly flowering-correlated genes left, for which BGS has expression levels characteristic of early flowering plants, are functionally diverse and do not include any known flowering time genes^33^.

Analysis of expression similarities between BGS and railway vs mountain expression patterns were performed using a custom-built metric on the results of two t-tests: *test* 1 compares the expression levels of railways versus mountains while grouping BGS with the mountain and *test* 2 while grouping BGS with the railways. The p-values of these two tests were then corrected for false discovery rate and their log-ratio computed for every gene differentially expressed between railways and mountains (2 sample t-test). We thus obtained a statistic called (RW/MT_*stat*_; Eq. 1), that was positive when the expression of a gene in BGS was closer to that seen in railway populations (*pval* 1 > *pval* 2) and negative when BGS levels were more similar to mountains.

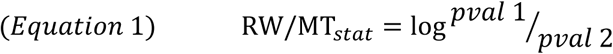

### Differentiation analysis

To test for genetic differentiation, we used our previously published genomic short read sequences for *A*. *arenosa*^40,67^ that we complemented with similarly processed genomes to reach 6 TBG, 8 STE, 8 BGS, 10 GU, 7 HO and 8 KA individuals for a total of 47 individuals over 6 populations. We aligned reads to the *A*. *lyrata* genome^32^ using BWA^68^ and re-aligned unmapped reads using Stampy^63^. We calculated F_ST_^69^ and Fay and Wu’s H^45^ and performed PCA analysis after genotyping the alignments with GATK^70^ only considering bi-allelic sites with a sequencing depth per individual of 4 or more (2.9 million SNPs).

For population structure analyses, we used STRUCTURE (Pritchard et al.) version 2.3.4 on 627 016 SNPs with a sequencing depth per individual of 8 or more (increased for computing memory purposes) with K values (number of groupings) ranging from 1 to 6.

Both Patterson’s *D*-statistic and modified *f*-statistics (*f̂_hom_* and *f̂_d_*) were calculated as described by Martin et al.^44^ using the frequency of derived alleles at each site in each population instead of binary counts:

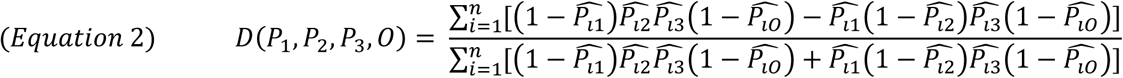

where *P*_1_, *P*_2_, *P*_3_ are the three populations used as background, receiver, and donor and *O* is *A*. *lyrata* (Fig. 3C). *p̂_lj_*are the observed frequencies of SNP *i* in population *P_j_*. As we polarized alleles frequencies in our sample using a panel of 24 *A*. *lyrata* genomes, we simplified *Equation 2* with *p̂_lO_*=0. Similarly, we calculated the modified *f*-statistic *f̂_hom_* described in Martin et al.^44^ as:

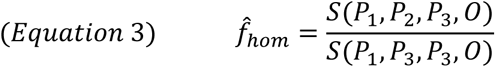

where *S*(*P*_1_, *P*_2_, *P*_3_, *0*) represents the numerator of *Equation 2*. We then split the genome into blocks of 50-kb-long which is greater than the very quick decay LD observed in *A*. *arenosa* in order to avoid correlation between blocks. We used a leave-one-out jackknife approach on these blocks to evaluate the confidence intervals of our genome-wide estimates (*D* and *f̂_hom_*). To identify candidate introgression loci on a gene scale, we used the modified *f*-statistic *f̂_d_* as:

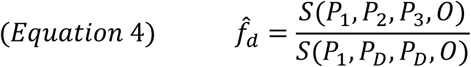

where *P_D_* is the highest value among *P*_2_ and *P*_3_ in order to take into account potential bidirectional gene-flow and incomplete lineage sorting which could give on a small window basis values of *f̂_hom_* negative or above 1. We then calculated *f̂_d_* for each gene when more than 25 informative SNPs were found across the gene annotation. We selected genes with high *f̂_d_* values for each of the four railway-mountain couples (*f̂_d_*(*P*_1_,*P*_2_,*P*_3_) > 3 ∗ *f̂_hom_*(*P*_1_,*P*_2_,*P*_3_)) in order to also take into account windows where introgression is high but introgressed alleles have not yet reached their equilibrium frequencies in BGS.

For graphic representation purposes (Fig. 6B), gene-wise estimates of G_ST_, Fay & Wu’s H, and Tajima’s D were obtained using the most extreme value of all windows overlapping a gene annotation. We then used the least extreme values of gene-wise G_ST_ calculated between the four railway-mountain couples (STE-HO, STE-KA, TBG-HO, and TBG-KA) as estimates of railway-mountain differentiation, the least extreme values of gene-wise Fay & Wu’s H and Tajima’s D in STE and TBG, to plot along the least extreme gene-wise *f̂_d_* estimates obtained for each of the four railway-mountain couples.

### Cloning and transgenic approach

We synthesized single strand cDNA from 500ng of total RNA of BGS and KA and PCR-amplified both *AaFLC* with primers 5’-CCCTCTCGGAGACAGAAGCCATGG-3’ (forward) and 5’-AGGTGGCTAATTAAGCAGCGGGAGAGTCAC-3’ (reverse). The PCR products were then introduced in pBluescript and sequenced using bounding M13 primers. The transgenes selected for transgenic confirmation were then cloned into pGREEN^71^ with the CMV 35S promoter and the Rbsc terminator. All constructs were then transformed using *Agrobacterium tumefaciens*, strain GV3101 by floral dipping^72^ into the early-flowering background *A*. *thaliana* line Col-0 *flc*-*3* obtained from R. Amasino, which we phenotyped for flowering time measured as leaf number at bolting (LNB) as well as days to bolt in our growth conditions (Fig. 5A). T1 seeds were selected on kanamycin (50 ug/ml) MS plates and resistant seedlings were transferred after a week to soil where they were phenotyped for flowering time (LNB and days to bolt). Leaf-tissue from all T1 plants was collected at 3-weeks and transgene expression was quantified by q-PCR and plotted against the flowering-time of each individual to take into account potential silencing effects of the transgene (variegation and gene-silencing). LNB showed the strongest correlation with transgene expression overall and was then used as the main proxy for flowering time. q-PCR was carried out on a Stratagene Mx3005P machine (Stratagene) with an annealing temperature of 55°C by using Taq DNA polymerase (New-England BioLabs). Reactions were carried out in triplicate, and we normalized *FLC* expression against expression of *ACTIN* using the 2^−□□CT^ method taking into account each primer’s efficiency as described in BIO-RAD Real Time PCR Applications Guide. The standard deviation of each biological replicate was calculated using a first order propagation of error formula on the variance of the technical replicates. We used cDNA-specific primers 5’-CAGCTTCTCCTCCGGCGATAACCTGG-3’ and 5’-GGCTCTGGTTACGGAGAGGGCA-3’ for *FLC* (87% efficiency) and 5’-CGTACAACCGGTATTGTGCTGGAT-3’ and 5’-ACAATTTCCCGCTCTGCTGTTGTG-3’ for *ACT* (91% efficiency).

### Accession numbers

RNAseq read data have been deposited in the NCBI SRA database under accession number SRP###### within the NCBI BioProject PRJNA######.

**Figure S1. Differential expression between railway and mountain accessions** Volcano plots of differential expression (q-value) against log expression ratios between railway and mountain accessions (excluding BGS) within whole transcriptome. 5% most differentially expressed (two-tailed log-ratio) are highlighted in green and within these, flowering-time genes (FT) are in red.

**Figure S2. Paralogue-specific *FLC* expression** Relative expression of *AaFLC1* (light grey) and *AaFLC2* (dark grey) across mountain populations and BGS.

**Figure S3. Correlation between flowering time and transgene expression** (A, B) Correlations between flowering time, measured as leaves number at bolting (LNB), and relative *FLC* expression in transgenic T1 lines for *AaFLC1* and *AaFLC2* 35S-driven cDNA transgenes of KA (A) and BGS (B). The regression line is represented in dotted line surrounded by the confidence intervals (shaded area). Black triangles mark the two late-flowering individuals obtained with BGS *AaFLC1* transgenes.

**Figure S4. Phenotypic distribution of flowering time and PVE distribution across the *FLC*-*CO* region in TBG × SWA and BGS × KA F_2_s** (A, B) Distribution of flowering time (Days to Bolting) in phenotyped (grey) and sequenced (blue) in TBG × SWA (A) and BGS × KA (B) F2 individuals. (C) Distribution of percentages of variance explained (PVE) across the *FLC*-*CO* region in TBG × SWA and BGS × KA. PVE distributions are shown for each cross above the gene models for the region. Single marker model (SMM) percent variance explained (PVE) are plotted in grey on the primary (left) y-axis, while semi-partial correlation coefficients (SPC) from the multiple linear model are in blue against the secondary y-axis.

**Figure S5. Patterns of differential expression over FT-peak region** (A) Principal Component Analysis (PCA) of gene expression levels within FT-peak region. (B) Volcano plots of differential expression (q-value) against log expression ratios between railway and mountain accessions (excluding BGS) within FT-peak region. 5% most differentially expressed (two-tailed log-ratio) are highlighted in green and within these, flowering-time genes (FT) are in red.

**Figure S6. Marks of railway-specific selection on *CONSTANS*** (A) Marks of differentiation between one railway population (TBG) and two mountain populations (HO and KA) evaluated with G_ST_ across *CO* region. Dotted lines are respective genome-wide 1% threshold levels. (B) Marks of positive selection in railway populations TBG, STE, and BGS measured by Fay and Wu’s H on 200kb region surrounding CO, with genome-wide 5% threshold levels (dotted lines).

**Figure S7. Rail networks in central Europe from 1849-1861.** Map showing the rail network in Germany and surrounding areas from 1849. The railways are indicated as solid bold black lines. Lines added by 1861 are shown as dotted lines illustrating the rapid expansion of a widely connected transport network. Map image is public domain and obtained from Wikipedia: https://en.wikipedia.org/wiki/History_of_rail_transport_in_Germany.

## REFERENCES

1. Hall, M. C. & Willis, J. H. Divergent selection on flowering time contributes to local adaptation in Mimulus guttatus populations. Evolution (N. Y). 60, 2466–2477 (2006).

2. Franks, S. J., Sim, S. & Weis, A. E. Rapid evolution of flowering time by an annual plant in response to a climate fluctuation. Proc. Natl. Acad. Sci. U. S. A. 104, 1278–82 (2007).

3. Fox, G. A. Drought and the evolution of flowering time in desert annuals. American Journal of Botany 77, 1508–1518 (1990).

4. Sherrard, M. E. & Maherali, H. The adaptive significance of drought escape in Avena barbata, an annual grass. Evolution (N. Y). 60, 2478–2489 (2006).

5. Mckay, J. K., Richards, J. H. & Mitchell-Olds, T. Genetics of drought adaptation in Arabidopsis thaliana: I. Pleiotropy contributes to genetic correlations among ecological traits. Mol. Ecol. 12, 1137–1151 (2003).

6. Baker, H. G. The Evolution of Weeds. Annu. Rev. Ecol. Syst. 5, 1–24 (1974).

7. Wu, C. A., Lowry, D. B., Nutter, L. I. & Willis, J. H. Natural variation for drought-response traits in the Mimulus guttatus species complex. Oecologia 162, 23–33 (2010).

8. Schweinsberg, F., Abke, W., Rieth, K., Rohmann, U. & Zullei-Seibert, N. Herbicide use on railway tracks for safety reasons in Germany? Toxicol. Lett. 107, 201–205 (1999).

9. Hartl, D. & Clark, A. Principles of Population Genetics. (Sinauer Associates, 1998).

10. Weinig, C. Rapid Evolutionary Responses to Selection in Heterogeneous Environments among Agricultural and Nonagricultural Weeds. Int. J. Plant Sci. 166, 641–647 (2005).

11. Kent, D. H. Senecio squalidus L. in the British Isles-2, the spread from Oxford (1879–1939). Proc. Bot. Soc. Br. Isles 3, 375–379 (1960).

12. Mack, R. N. Alien Plant Invasion into the Intermountain West: A Case History. (Springer New York, 1986). doi:10.1007/978-1-4612-4988-7_12

13. Matlack, G. Exotic plant species in Mississippi, USA: Critical issues in management and research. Nat. AREAS J. 22, 241–247 (2002).

14. Christen, D. & Matlack, G. The Role of Roadsides in Plant Invasions: a Demographic Approach. Conserv. Biol. 20, 385–391 (2006).

15. Clauss, M. J. & Koch, M. A. Poorly known relatives of Arabidopsis thaliana. Trends Plant Sci. 11, 449–59 (2006).

16. O’Kane, S. L. A Synopsis of Arabidopsis (Brassicaceae). Novon 7, 323–327 (1997).

17. Schmickl, R., Paule, J., Klein, J., Marhold, K. & Koch, M. A. The evolutionary history of the Arabidopsis arenosa complex: diverse tetraploids mask the Western Carpathian center of species and genetic diversity. PLoS One 7, e42691 (2012).

18. Arnold, B., Kim, S.-T. & Bomblies, K. Single Geographic Origin of a Widespread Autotetraploid Arabidopsis arenosa Lineage Followed by Interploidy Admixture. Mol. Biol. Evol. 32, 1382–95 (2015).

19. Baduel, P., Arnold, B., Weisman, C. M., Hunter, B. & Bomblies, K. Habitat-Associated Life History and Stress-Tolerance Variation in *Arabidopsis arenosa*. Plant Physiol. 171, 437–451 (2016).

20. Searle, I. et al. The transcription factor FLC confers a flowering response to vernalization by repressing meristem competence and systemic signaling in Arabidopsis. Genes Dev. 20, 898–912 (2006).

21. Andrés, F. & Coupland, G. The genetic basis of flowering responses to seasonal cues. Nat. Rev. Genet. 13, 627–39 (2012).

22. Michaels, S. D. & Amasino, R. M. FLOWERING LOCUS C Encodes a Novel MADS Domain Protein That Acts as a Repressor of Flowering. Plant Cell 11, 949–956 (1999).

23. Gazzani, S., Gendall, A. R., Lister, C. & Dean, C. Analysis of the molecular basis of flowering time variation in Arabidopsis accessions. Plant Physiol. 132, 1107–14 (2003).

24. Lempe, J. et al. Diversity of flowering responses in wild Arabidopsis thaliana strains. PLoS Genet. 1, 109–18 (2005).

25. Shindo, C. et al. Role of FRIGIDA and FLOWERING LOCUS C in determining variation in flowering time of Arabidopsis. Plant Physiol. 138, 1163–73 (2005).

26. Werner, J. D. et al. FRIGIDA-independent variation in flowering time of natural Arabidopsis thaliana accessions. Genetics 170, 1197–207 (2005).

27. Méndez-Vigo, B., Picó, F. X., Ramiro, M., Martínez-Zapater, J. M. & Alonso-Blanco, C. Altitudinal and climatic adaptation is mediated by flowering traits and FRI, FLC, and PHYC genes in Arabidopsis. Plant Physiol. 157, 1942–55 (2011).

28. Salomé, P. A. et al. Genetic architecture of flowering-time variation in Arabidopsis thaliana. Genetics 188, 421–33 (2011).

29. Méndez-Vigo, B. et al. Environmental and genetic interactions reveal FLC as a modulator of the natural variation for the plasticity of flowering in Arabidopsis. Plant. Cell Environ. (2015). doi: 10.1111/pce.12608

30. Wang, R. et al. PEP1 regulates perennial flowering in Arabis alpina. Nature 459, 423–7 (2009).

31. Albani, M. C. et al. PEP1 of Arabis alpina is encoded by two overlapping genes that contribute to natural genetic variation in perennial flowering. PLoS Genet. 8, e1003130 (2012).

32. Hu, T. T. et al. The Arabidopsis lyrata genome sequence and the basis of rapid genome size change. Nat. Genet. 43, 476–81 (2011).

33. Fornara, F., de Montaigu, A. & Coupland, G. SnapShot: Control of Flowering in Arabidopsis. Cell 141, 550–550 (2010).

34. Li, D. et al. A repressor complex governs the integration of flowering signals in Arabidopsis. Dev. Cell 15, 110–20 (2008).

35. Michaels, S. D. Loss of FLOWERING LOCUS C Activity Eliminates the Late-Flowering Phenotype of FRIGIDA and Autonomous Pathway Mutations but Not Responsiveness to Vernalization. PLANT CELL ONLINE 13, 935–942 (2001).

36. Jung, J.-H., Ju, Y., Seo, P. J., Lee, J.-H. & Park, C.-M. The SOC1-SPL module integrates photoperiod and gibberellic acid signals to control flowering time in Arabidopsis. Plant J. 69, 577–588 (2012).

37. Samach, A. et al. Distinct roles of CONSTANS target genes in reproductive development of Arabidopsis. Science 288, 1613–6 (2000).

38. Yoo, S. K. et al. CONSTANS Activates SUPPRESSOR OF OVEREXPRESSION OF CONSTANS 1 through FLOWERING LOCUS T to Promote Flowering in Arabidopsis. PLANT Physiol. 139, 770–778 (2005).

39. Nah, G. & Jeffrey Chen, Z. Tandem duplication of the FLC locus and the origin of a new gene in Arabidopsis related species and their functional implications in allopolyploids. New Phytol. 186, 228–38 (2010).

40. Hollister, J. D. et al. Genetic adaptation associated with genome-doubling in autotetraploid Arabidopsis arenosa. PLoS Genet. 8, e1003093 (2012).

41. Arnold, B. J. et al. Borrowed alleles and convergence in serpentine adaptation. Proc. Natl. Acad. Sci. 113, 8320–8325 (2016).

42. Pritchard, J. K. et al. Inference of population structure using multilocus genotype data. Genetics 155, 945–59 (2000).

43. Evanno, G., Regnaut, S. & Goudet, J. Detecting the number of clusters of individuals using the software structure: a simulation study. Mol. Ecol. 14, 2611–2620 (2005).

44. Martin, S. H., Davey, J. W. & Jiggins, C. D. Evaluating the Use of ABBA-BABA Statistics to Locate Introgressed Loci. Mol. Biol. Evol. 32, 244–257 (2015).

45. Fay, J. C. & Wu, C.-I. Hitchhiking Under Positive Darwinian Selection. Genetics 155, 1405–1413 (2000).

46. Nei, M. Analysis of gene diversity in subdivided populations. Proc. Natl. Acad. Sci. U. S. A. 70, 3321–3 (1973).

47. Tajima, F. Statistical method for testing the neutral mutation hypothesis by DNA polymorphism. Genetics 123, 585–95 (1989).

48. Kolář, F. et al. Northern glacial refugia and altitudinal niche divergence shape genome-wide differentiation in the emerging plant model *Arabidopsis arenosa*. Mol. Ecol. 25, 3929–3949 (2016).

49. Whitney, K. D. et al. Quantitative trait locus mapping identifies candidate alleles involved in adaptive introgression and range expansion in a wild sunflower. Mol. Ecol. 24, 2194–2211 (2015).

50. Whitney, K. D., Randell, R. A. & Rieseberg, L. H. Adaptive introgression of abiotic tolerance traits in the sunflower Helianthus annuus. New Phytol. 187, 230–239 (2010).

51. Whitney, K. D., Randell, R. A. & Rieseberg, L. H. Adaptive Introgression of Herbivore Resistance Traits in the Weedy Sunflower *Helianthus annuus*. Am. Nat. 167, 794–807 (2006).

52. Song, Y. et al. Adaptive Introgression of Anticoagulant Rodent Poison Resistance by Hybridization between Old World Mice. Current Biology 21, (2011).

53. Hedrick, P. W. Adaptive introgression in animals: examples and comparison to new mutation and standing variation as sources of adaptive variation. Mol. Ecol. 22, 4606–4618 (2013).

54. Castric, V., Bechsgaard, J., Schierup, M. H. & Vekemans, X. Repeated Adaptive Introgression at a Gene under Multiallelic Balancing Selection. PLoS Genet. 4, e1000168 (2008).

55. Hepworth, S. R. Antagonistic regulation of flowering-time gene SOC1 by CONSTANS and FLC via separate promoter motifs. EMBO J. 21, 4327–4337 (2002).

56. Yano, M. et al. Hd1, a Major Photoperiod Sensitivity Quantitative Trait Locus in Rice, Is Closely Related to the Arabidopsis Flowering Time Gene CONSTANS. PLANT CELL ONLINE 12, 2473–2484 (2000).

57. Österberg, M. K. et al. Naturally Occurring Indel Variation in the Brassica nigra COL1 Gene Is Associated With Variation in Flowering Time. Genetics 161, 749–764 (2002).

58. Rosas, U. et al. Variation in Arabidopsis flowering time associated with cis-regulatory variation in CONSTANS. Nat. Commun. 5, (2014).

59. Takahashi, Y. & Shimamoto, K. Heading date 1 (Hd1), an ortholog of Arabidopsis CONSTANS, is a possible target of human selection during domestication to diversify flowering times of cultivated rice. Genes Genet. Syst. 86, 175–182 (2011).

60. Huerta-Sánchez, E. et al. Altitude adaptation in Tibetans caused by introgression of Denisovan-like DNA. Nature 512, 194–197 (2014).

61. Jørgensen, M. H., Ehrich, D., Schmickl, R., Koch, M. A. & Brysting, A. K. Interspecific and interploidal gene flow in Central European Arabidopsis (Brassicaceae). BMC Evol. Biol. 11, 346 (2011).

62. Kim, D. et al. TopHat2: accurate alignment of transcriptomes in the presence of insertions, deletions and gene fusions. Genome Biol. 14, R36 (2013).

63. Lunter, G. & Goodson, M. Stampy: A statistical algorithm for sensitive and fast mapping of Illumina sequence reads. Genome Res. 21, 936–939 (2011).

64. Anders, S. HTSeq: Analysing high-throughput sequencing data with Python. (2011).

65. Anders, S. & Huber, W. Differential expression analysis for sequence count data. Genome Biol. 11, 1–12 (2010).

66. Wang, J., Tian, L., Lee, H.-S. & Chen, Z. J. Nonadditive regulation of FRI and FLC loci mediates flowering-time variation in Arabidopsis allopolyploids. Genetics 173, 965–74 (2006).

67. Yant, L. et al. Meiotic adaptation to genome duplication in Arabidopsis arenosa. Curr. Biol. 23, 2151–6 (2013).

68. Li, H. & Durbin, R. Fast and accurate short read alignment with Burrows–Wheeler transform. Bioinformatics 25, 1754–1760 (2009).

69. Weir, B. S. Genetic data analysis. Methods for discrete population genetic data. (Sinauer Associates, Inc. Publishers, 1990).

70. McKenna, A. et al. The Genome Analysis Toolkit: a MapReduce framework for analyzing next-generation DNA sequencing data. Genome Res. 20, 1297–303 (2010).

71. Hellens, R. P., Edwards, E. A., Leyland, N. R., Bean, S. & Mullineaux, P. M. pGreen: a versatile and flexible binary Ti vector for Agrobacterium-mediated plant transformation. Plant Mol. Biol. 42, 819–832 (2000).

72. Ellis, J. R. in Plant Molecular Biology LABFAX 253–279 (Bios Scientific Publishers, 1993).

